# A ratiometric biochemical framework reveals strain-specific metabolic allocation strategies in brook trout liver

**DOI:** 10.64898/2026.08.09.743818

**Authors:** Katie A. Edwards, Eileen A. Randall, Clifford E. Kraft, Banshika M. Mangal, Dennis Kleiner

## Abstract

Brook trout (*Salvelinus fontinalis*) exhibit strain-level variation in growth performance, environmental tolerance, and survival, yet the biochemical mechanisms underlying these differences remain poorly understood. We developed and applied a ratiometric biochemical framework integrating the pentose-phosphate pathway (PPP) and glutathione metabolism to characterize strain-specific hepatic metabolic organization in brook trout. Five strains reared under standardized conditions differed significantly in hepatic soluble protein density, glutathione pool size, total NADP(H) concentration, and activities of glucose-6-phosphate dehydrogenase (G6PDH), glutathione reductase (GR), and transketolase (TKT). These differences were not uniformly coordinated across pathways, demonstrating that metabolic phenotype cannot be inferred from individual biomarkers alone. Derived ratios describing oxidative-to-non-oxidative PPP capacity (G6PDH/TKT) and glutathione buffering relative to recycling capacity ((GSH+GSSG)/GR) resolved distinct patterns of metabolic allocation among strains. Despite shared ancestry, the Temiscamie (TEM) strain and its domestic × TEM hybrid (TXD) exhibited markedly divergent metabolic phenotypes, demonstrating that closely related strains can differ substantially in hepatic metabolic organization. Together, these findings identify relative allocation among interconnected metabolic pathways as an axis of physiologic diversity and establish a ratiometric approach for comparing metabolic organization across populations and species.

**Graphical abstract:** Hepatic metabolic phenotypes of brook trout strains were characterized by integrating pentose phosphate pathway enzyme capacities, glutathione metabolism, NADP(H) availability, and soluble protein into a ratiometric framework. Ratios distinguish investment in oxidative versus non-oxidative PPP capacity (G6PDH/TKT), antioxidant buffering versus glutathione recycling capacity (total glutathione/GR), and hepatic protein density (soluble protein/liver mass), revealing distinct metabolic organization among strains.

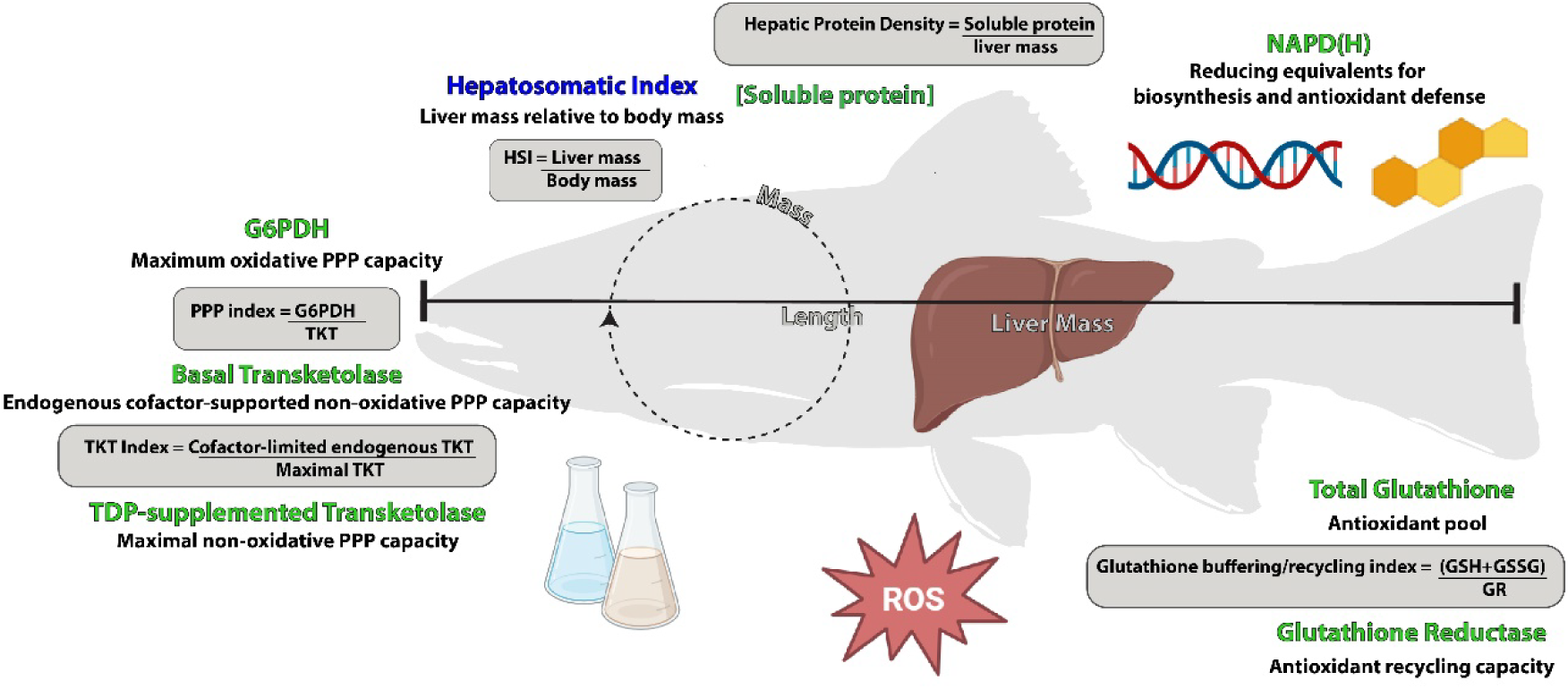

**Highlights:**

- A ratiometric framework was developed to characterize hepatic metabolic organization in brook trout
- Glutathione buffering and recycling capacity distinguish alternative redox phenotypes
- Investment in oxidative and non-oxidative PPP capacity varies independently among strains
- G6PDH/TKT and total glutathione (GSH+GSSG)/GR reveal distinct metabolic phenotypes
- Ratiometric indices provide a framework for interpreting redox metabolism and carbon allocation

## Introduction

Brook trout (*Salvelinus fontinalis*) inhabit aquatic ecosystems ranging from small streams to bodies of water as large as the Laurentian Great Lakes and coastal ocean habitats. Their native range spans the southern Appalachians through northern Quebec and the Great Lakes region, and brook trout have been widely introduced in cold-water rivers in western North America, Europe, Argentina, Japan and Australia.[1, 2] As a stenothermic salmonine fish, brook trout exhibit optimal growth and performance at temperatures below 18°C and experience sublethal stress above ∼20°C.[3] Although brook trout populations are often compared based on morphology, growth, survival, and habitat tolerance,[4–7] comparatively little is known about how genetically distinct strains differ in their baseline biochemical organization. In particular, the degree to which strains vary in their allocation of metabolic capacity toward redox balance, antioxidant buffering, and carbon routing under non-stressed conditions has not been considered with regard to their ability to thrive in a wide range of habitat conditions, even though baseline metabolic organization defines the constraints within which physiological plasticity must operate.

At the cellular level, redox balance is tightly linked to the organization of the pentose phosphate pathway (PPP) that provides reducing equivalents in the form of NADPH that transfers electrons in redox reactions, while interfacing dynamically with glycolysis and biosynthetic pathways (Fig. 1). The oxidative arm of the PPP, initiated by glucose-6-phosphate dehydrogenase (G6PDH), generates NADPH from NADP^+^, while converting glucose-6-phosphate (G6P) to ribulose-5-phosphate (Ru5P).[8–10] NADPH provides an essential reducing currency by providing electrons to support antioxidant defenses, lipid and cholesterol synthesis, xenobiotic metabolism, nitric oxide signaling, and folate metabolism.[11] In contrast, the non-oxidative arm of the PPP, catalyzed in part by transketolase (TKT), redistributes carbon skeletons to support nucleotide synthesis via ribose-5-phosphate (R5P) and glycolytic flux without directly generating or consuming reducing equivalents.[12, 13] G6PDH and TKT are the rate-limiting enzymes of the oxidative and non-oxidative phases of the PPP, respectively.[9, 13]

**Fig. 1.**
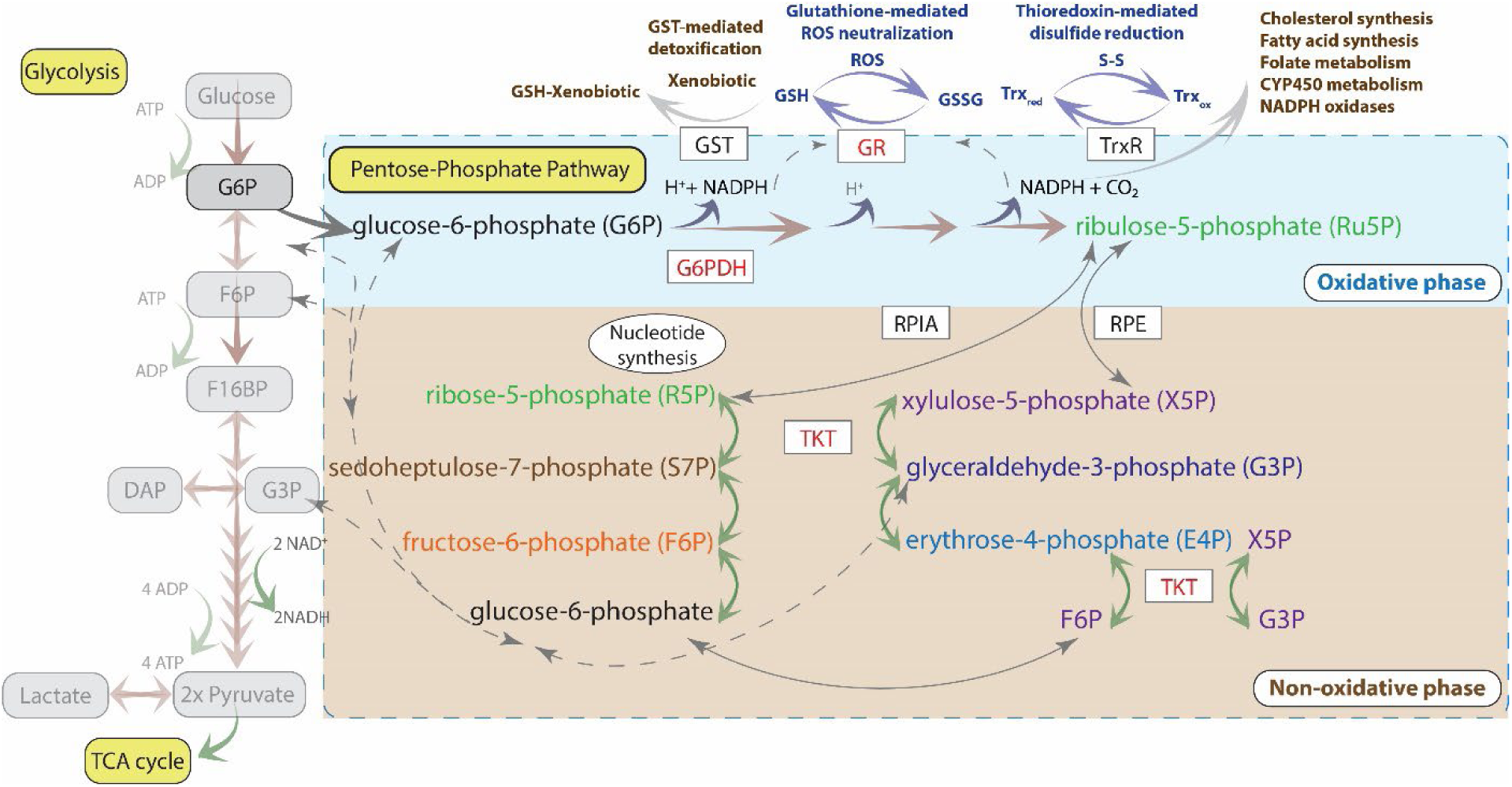
Integration of the pentose phosphate pathway (PPP) with glycolysis, glutathione-dependent xenobiotic metabolism, and antioxidant defense. The oxidative (blue) and non-oxidative (brown) phases of the PPP generate NADPH and interconvert glycolytic and pentose phosphate intermediates, respectively. NADPH produced by glucose-6-phosphate dehydrogenase (G6PD) supports glutathione reductase (GR), thioredoxin reductase (TrxR), glutathione-S-transferase (GST)-mediated detoxification, and other NADPH-dependent biosynthetic pathways. The enzymes assayed in this study (in red) were G6PDH, transketolase (TKT), and GR. Not all reactions in these pathways are explicitly shown.

The balance between oxidative and non-oxidative PPP activity reflects a fundamental metabolic divergence, allocating carbon toward redox support versus biosynthetic flexibility, with important implications for disease and responses to oxidative stress.[13–15] Ru5P can be synthesized catabolically during the oxidative phase and subsequently isomerized to form R5P and xylulose-5-phosphate (X5P) for DNA and RNA production in the non-oxidative phase, or R5P can be synthesized anabolically via reversal of non-oxidative-phase reactions using glycolytic intermediates.[10, 15] The oxidative PPP enables simultaneous production of NADPH upstream of R5P formation, thereby meeting the demand for reducing equivalents.[10, 16] However, we postulate that routing carbons directly from glycolysis through the non-oxidative phase may be preferable when carbon conservation is metabolically advantageous, as the oxidative phase loses carbon via CO_2_ release. The metabolic balance is further shaped by downstream redox systems, particularly the glutathione network. Glutathione reductase (GR) uses NADPH to regenerate reduced glutathione (GSH) from oxidized glutathione (GSSG).[17, 18] The size of the glutathione pool and its recycling capacity together determine redox resilience, while their relationship to NADPH availability reflects broad metabolic budgeting strategies.

The liver is a particularly informative tissue for examining these relationships. Hepatic metabolism integrates nutrient processing, gluconeogenesis, detoxification, and antioxidant defense, and exhibits strong temperature dependence in stenothermic fish.[19–21] Liver tissue also displays high antioxidant enzyme activities relative to muscle,[22] making it a central organ for maintaining redox homeostasis under routine metabolic demand. The liver is also the primary organ for glutathione synthesis, in part due to its unique ability to synthesize cysteine, the rate-limiting amino acid of the glutathione tripeptide.[23, 24]

Brook trout populations in northeastern North America exhibit persistent, watershed-specific genetic structure, indicating that contemporary strains retain signatures of geographic and evolutionary divergence despite long-term hatchery management.[25] In this study, we examined baseline hepatic metabolic organization across five brook trout strains of different origins: Assinica (ASN), Temiscamie (TEM), Temiscamie X Domestic F1 hybrid (TXD), Horn Lake (HRN), and Little Tupper (LT). ASN and TEM were brought to New York from Canada and have been maintained in a limited number of New York lakes for the past 70 years. HRN and LT strains originated in Adirondack lakes and have been maintained in a small number of lakes by the New York State Department of Environmental Conservation (DEC).[26–28] By contrast with the wild strains, TXD is a TEM and domestic cross widely raised and stocked throughout New York due to its hatchery growth, survival, tolerance of low-pH waters, and ease of capture.[28, 29] These strains differ in geographic origin, water preferences, management history, and selection pressures,[6, 30] allowing us to examine how metabolic allocation strategies diverge within a species of fish that inhabits wide-ranging environmental conditions. Using a combination of enzyme activities, metabolite pool measurements, and derived ratiometric indices, we quantified strain-specific differences across two interrelated metabolic axes:

1. Oxidative versus non-oxidative PPP capacity (G6PDH, total NADP(H), and basal/supplemented TKT)
2. Redox buffering versus recycling capacity (GR activity and glutathione pool size)

The study fish were reared under identical common hatchery conditions for eight months, allowing strain-level differences in hepatic metabolism to be evaluated while minimizing environmental variation. By establishing how metabolic systems of these fish are organized, this study provides a mechanistic framework to evaluate strain-specific metabolic allocation strategies in brook trout.

## Materials and Methods

### Materials

The following reagents were purchased from Sigma: α-GDH-TPI from rabbit muscle, glutathione reductase from Baker’s yeast (*Saccharomyces cerevisiae*), ribose-5-phosphate (R5P), thiamine diphosphate (TDP), and NADH. The following reagents and enzymes were purchased from Thermo Scientific: reduced glutathione (GSH), glucose-6-phosphate dehydrogenase (yeast), NADPH, 5-disulfosalicylic acid, and Micro BCA Protein Assay Kit. General supplies and buffer components were procured from Avantor or Fisher Scientific.

### Fish husbandry

Brook trout strains were raised from eggs by the following hatcheries in New York State: New Brandon Fisheries produced the Temiscamie, Temiscamie-Domestic F1 hybrids, Little Tupper, and Assinica strains and Warren County Fish Hatchery produced the Horn Lake strain. The fry/young fish were obtained in June of 2024 at an approximate length of ∼60 mm, then reared at the Little Moose field station (Old Forge, NY). Each strain was maintained in separate, identical 10-foot-diameter round concrete tanks under the same growing conditions in an enclosed pole barn setting for 8 months following arrival. The fish were fed BioVita Fry fish meal (BioOregon) by hand two to three times per day during normal business hours throughout the study period and the tanks were cleaned within two hours of feeding. In February of 2025, the individual fish included in this study were measured (total length, mm), weighed (g), and sacrificed in accordance with Cornell University IACUC procedure 2008-0009. Liver samples were extracted upon achieving postmortem status and immediately stored in liquid nitrogen for transport to the laboratory facility, where they were maintained at - 80°C until analysis.

### Protein extraction

∼200 mg of frozen liver tissue per sample was diced and weighed into 2 mL microcentrifuge tubes. Samples were kept on ice during weighing and returned immediately to -80°C storage. 3 mL/g of ice-cold 0.25 M sucrose was added to each tube, and the samples were homogenized with a hand-held homogenizer (Bio-Spec Tissue-Tearor™) at the lowest speed for 30 seconds while the tubes remained on ice. The homogenized samples were then centrifuged at 12,000 × g at 4°C for 20 min. before promptly aliquoting the supernatants and storing them at -80°C.

### Enzyme assays

#### Glucose-6-phosphate dehydrogenase (G6PDH) activity

G6PDH activity was assessed using procedures modified from Amcoff et al.[31] and adapted for use in a microplate format. The working reagent consisted of freshly prepared 1.150 mM glucose-6-phosphate (G6P) and 0.575 mM oxidized nicotinamide adenine dinucleotide phosphate (NADP^+^) in 100 mM Tris, pH 7.6, containing 0.004%(w/v) Tween-20. Liver extract supernatants were diluted 1:35 in Tris, pH 7.6. A sucrose-only blank was diluted similarly using the 0.25 M sucrose solution used for extraction. Then, 30 μL of the diluted samples and blanks were added in triplicate to a 96-well UV-transparent microtiter plate. 200 μL of the working reagent was added to all wells and immediately mixed. The wells were then read at 340 nm every 60 seconds for 1 hour, with 5 seconds of shaking between readings, with the plate reader equilibrated at 37°C. The plate was then read for pathlength correction at 975 nm and 900 nm.

#### Glutathione reductase (GR) activity

GR activity was assessed using procedures modified from Sigma-Aldrich[32] and adapted for use in a microplate format. The working reagent consisted of freshly prepared 1.150 mM oxidized glutathione (GSSG) and 0.115 mM reduced nicotinamide adenine dinucleotide phosphate (NADPH) in 200 mM potassium phosphate, pH 7.6, containing 2 mM EDTA and 0.004%(w/v) Tween-20. Liver extract supernatants were diluted 1:35 in Tris, pH 7.6. A sucrose-only blank was diluted similarly using the 0.25 M sucrose solution used for extraction. Then, 30 μL of the diluted samples and blanks were added in triplicate to a 96-well UV-transparent microtiter plate. 200 μL of the working reagent was added to all wells and immediately mixed. The wells were then read at 340 nm every 60 seconds for 1 hour, with 5 seconds of shaking between readings, with the plate reader equilibrated at 37°C. The plate was then read for pathlength correction at 975 nm and 900 nm.

#### Transketolase (TKT) activity

TKT activity was assessed following the procedures in Edwards *et al*,[33] except the samples were run more dilute (1:35) rather than 1:10 due to the greater TKT activity in brook trout.

#### Total NADP(H) (NADPH + NADP^+^)

The total NADP(H) concentration was determined using modifications of procedures in Wagner and Scott,[34] and adapted for use in a microplate format. Liver supernatants and a sucrose-only blank were diluted 1:45 in an extraction buffer composed of 20 mM nicotinamide, 100 mM sodium carbonate, and 20 mM sodium bicarbonate, then were frozen at -80°C overnight. The samples were gently thawed in a 22°C water bath, centrifuged at 16,000 × g at 4°C, and stored on ice. NADPH standards were prepared from 40 nM to 2.5 µM in the extraction buffer, and 100 µL of NADPH standards and 100 µL of samples were added in triplicate to a Corning Costar 3795 microtiter plate. A working reagent containing 1.6 mM phenazine ethosulfate (PES), 0.4 mM methylthiazolyldiphenyl-tetrazolium bromide (MTT), and 1 U/mL G6PDH in cycling buffer (composed of 177.5 mM Tris and 8.9 mM EDTA, pH 8.0) was freshly prepared. 100 µL of the working reagent was added to all wells. The plate was sealed and incubated at 37°C for 5 min. in the dark. 22 µL of 10.1 mM G6P in cycling buffer was prepared fresh and added to the wells immediately before shaking and reading at 570 nm every 60 seconds for 1 hour, with 5 seconds of shaking between readings, with the plate reader equilibrated at 37°C. The plate was then read for pathlength correction at 975 nm and 900 nm.

#### Total glutathione (GSH+GSSG)

The total glutathione concentration was determined using modifications of procedures described by Akerboom and Sies[35], further detailed in Sigma[36], and scaled for use in a microplate format. 45 µL of ice-cold 5% (w/v) sulfosalicylic acid (SSA) in water was added to 5 µL of the neat supernatants. The mixtures were vortexed, briefly centrifuged, and stored at 4°C for 10 minutes to ensure complete protein precipitation. The mixtures were centrifuged for 5 min. at 10,000 × g, then 10 µL of the supernatants were added in triplicate to a clear Corning Costar 3795 microtiter plate. 10 µL of reduced glutathione standards (52 μM to 200 µM) prepared in 5% (w/v) SSA were used in triplicate. A working reagent was freshly prepared, composed of 30 µg/mL 5,5′-dithiobis(2-nitrobenzoic acid) (DTNB) and 118 mU/mL glutathione reductase in 100 mM potassium phosphate, pH 7.0, containing 1 mM EDTA. 100 µL of the working reagent was added to all wells, then the plate was sealed and shaken in the dark at 350 rpm on a microplate mixer. 50 µL of 213 µM NADPH in 100 mM potassium phosphate, pH 7.0, containing 1 mM EDTA was then added. The plate was immediately shaken and read at 412 nm every 30 seconds for 10 min, with 5 seconds of shaking between reads, while the plate reader was equilibrated at 25°C. The plate was then read for pathlength correction at 975 nm and 900 nm.

#### Protein concentration assays

The Micro BCA assay was carried out on the same diluted extracts used for the samples, using bovine serum albumin (BSA) standards, following the manufacturer’s instructions.

#### Data analysis

The hepatosomatic index (HSI) was derived by dividing the liver mass by the body mass and expressing it as a percentage. For all analyses, wet liver tissue masses (not dehydrated) were used. All spectrophotometric data were pathlength-corrected using absorbance values at 900 nm and 975 nm and a K-factor of 0.173.[37, 38] The pathlength-corrected data were then normalized by deducting their initial OD value, and the maximum slopes of the OD versus time plots (minutes) were determined using GraphPad Prism, Version 10.2. The slopes for TKT, G6PDH, and GR were converted to pmol/min. We confirmed that the fish liver matrix at the maximum concentration used in the assays (1:35, 1.24 mg tissue/mL) did not interfere with the detection of NADPH at 340 nm. The concentration of total NADP(H) (NADPH + NADP^+^) in the original extracts was determined against the NADPH calibration curve in μM and accounted for the 1:45 sample dilution. To convert to pmol NADP(H)/mg tissue, the determined μmol/L concentration was multiplied by 3 to account for the original extract dilution (3 mL/g tissue) and unit conversions. The concentration of total GSH+GSSG in the original extracts was determined using the GSH calibration curve (mM) and accounted for the 1:10 sample dilution. To convert to pmol (GSH+GSSG)/mg tissue, the determined mmol/L concentration was multiplied by 3,000 to account for the original extract dilution (3 mL/g tissue) and unit conversions. In other assays, the data were then normalized to mg tissue by dividing by the respective masses within the wells. Although total protein masses were available for each assay, normalization was done throughout on a tissue mass basis.

#### Statistics

Means ± standard errors (SE) were calculated for each measured and derived parameter. Differences between strains were assessed by one-way ANOVAs with Tukey’s post hoc multiple comparison tests. Statistical significance was defined as p<0.05. The heatmap was generated for each strain by calculating Z-scores for each parameter across strains. To evaluate whether biochemical parameters varied among strains independently of body size, analyses of covariance (ANCOVAs) were performed with strain as a fixed factor and either body length or mass as a continuous covariate using the Standard Least Squares models platform in JMP. Plots and statistical analyses were carried out in JMP Student Edition (version 18.2.2) and GraphPad (version 11.0.2). Figures were drawn in Adobe Illustrator (version 30.3.)

## Results

Although strain differences are often characterized on a physiological, behavioral, or genetic basis, we present a framework for evaluating strain-specific metabolic differences at a biochemical level. We designed the study to measure enzyme activities (glucose-6-phosphate dehydrogenase (G6PDH), transketolase (TKT), and glutathione reductase (GR)) alongside key cellular redox currencies (NADP(H) and glutathione) to evaluate the relationship between maximal enzyme capacity and the availability of metabolites that support antioxidant defense, biosynthesis, and detoxification. We included G6PDH and TKT, as they are rate-limiting enzymes of the oxidative and non-oxidative PPP, respectively, to assess the relative investment in these branches of this pathway. NADP(H) was included to assess the pool of reducing equivalent currency available to support reduced glutathione regeneration, fatty acid and cholesterol biosynthesis, CYP450 metabolism, and folate metabolism. GR was included as an index of glutathione recycling capacity, which maintains the availability of reduced glutathione required to mediate ROS scavenging and detoxify xenobiotics. Measurements of the enzymes under substrate-saturating conditions reflect their maximal capacity and therefore their abundance and physiological limits rather than instantaneous metabolic flux. NADP(H) and total glutathione were assessed as their total available pools (NADP^+^ + NADPH and GSH + GSSG) for biochemical processes rather than their individual redox states. Evaluating these parameters independently reflects pathway capacity, while profiling them using ratiometric analyses indicates strain-specific metabolic strategies required under contrasting environmental conditions.

### Analysis of strain-dependence

The mean and standard error for each parameter, including TKT (basal and TDP-supplemented), G6PDH, and GR activities; total NADPH/NADP+; total glutathione (GSH + GSSG); extractable protein concentration; liver mass; length; and weight, and derived ratiometric parameters are presented in Table 1 on a per-strain basis. One-way ANOVAs with Tukey’s multiple-comparisons post hoc tests were conducted to assess differences among strains. Under substrate-saturating conditions (R5P for TKT, G6P for G6PDH, GSSG for GR), measured activities reflect strain-specific maximal enzyme capacity governed by expression and post-translational state, rather than in vivo substrate or redox limitation. This is consistent with differences in metabolic strategy, rather than instantaneous metabolic flux.

**Table 1.**
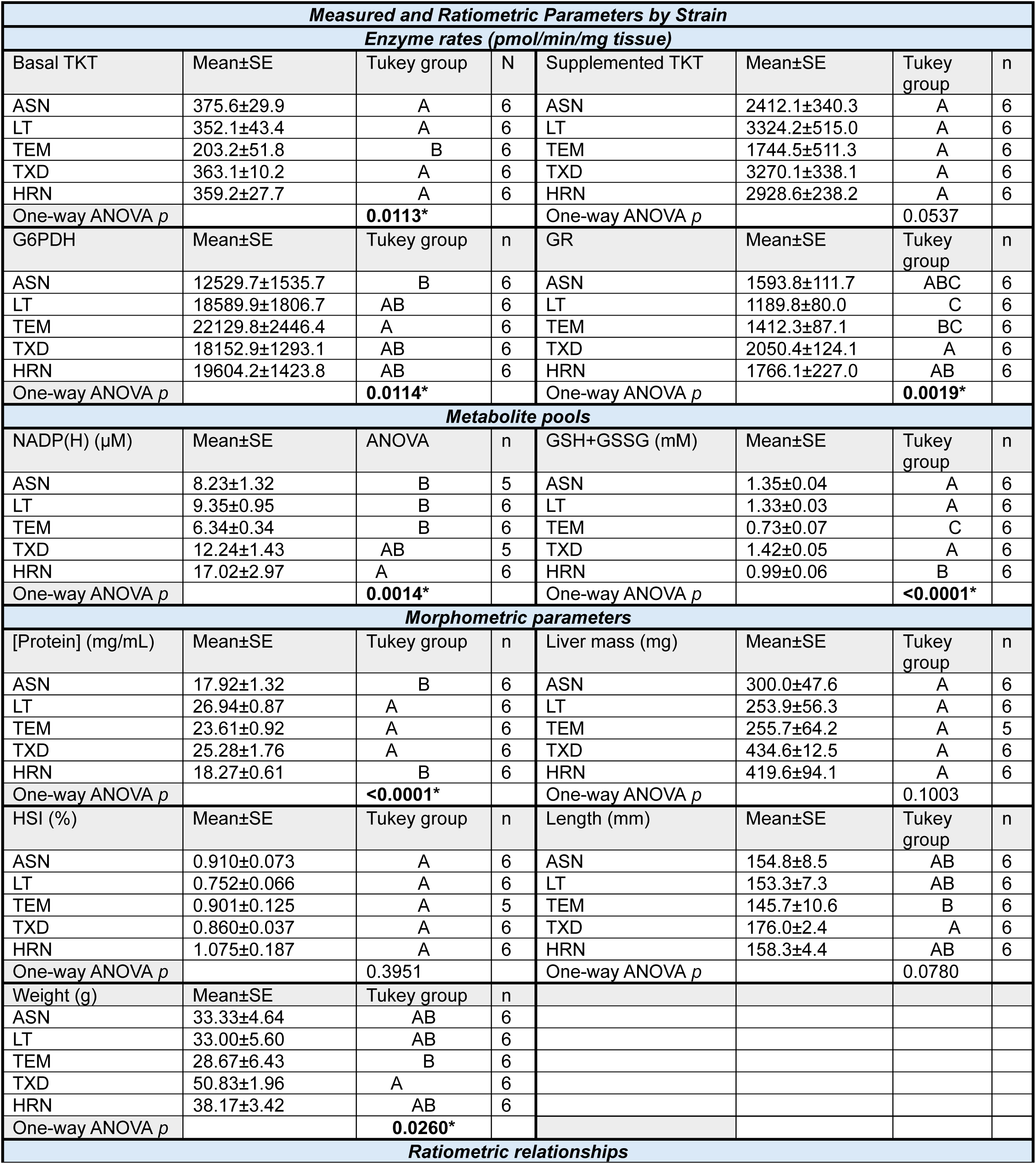
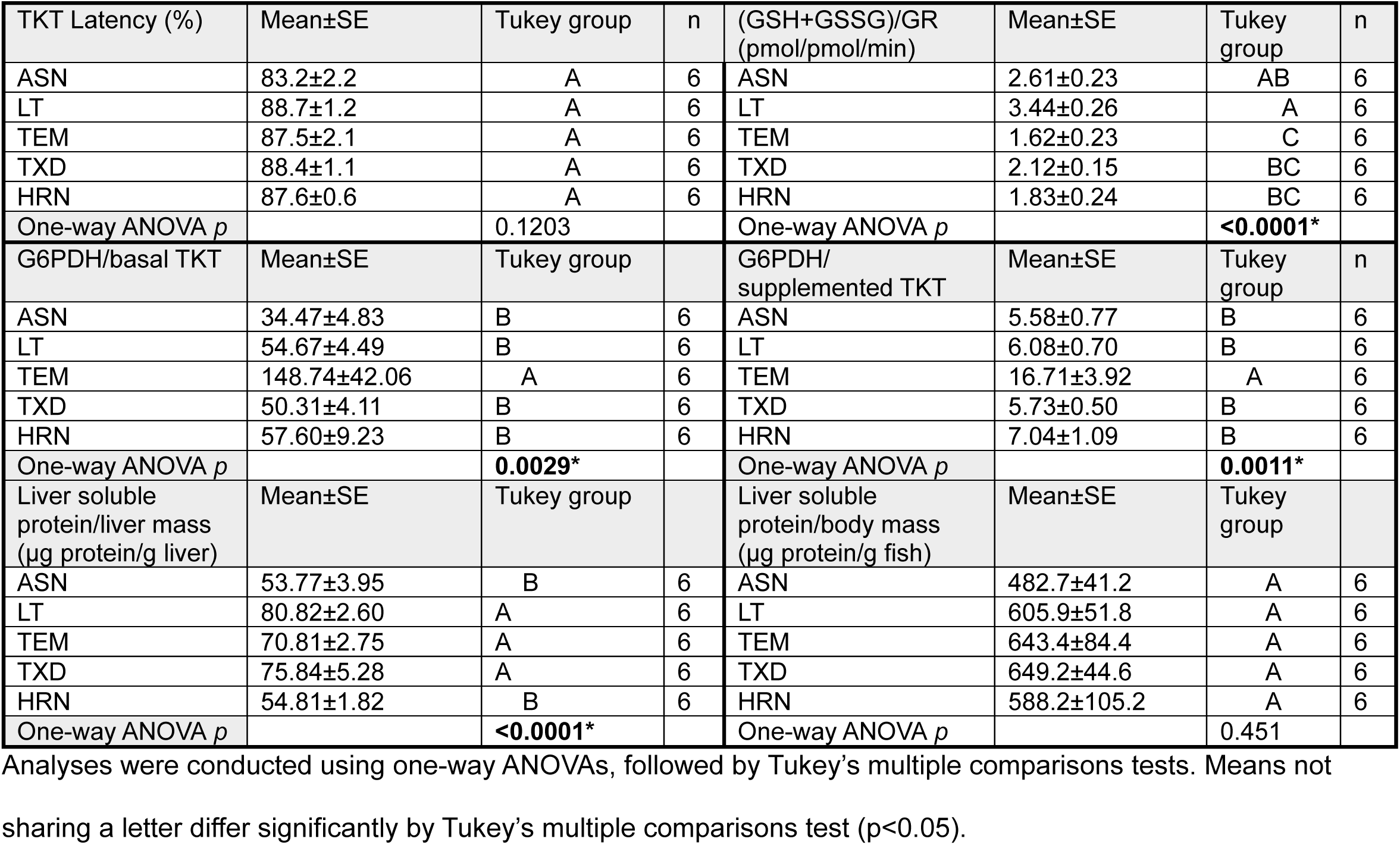
Enzyme, metabolite, morphometric, and ratiometric parameters.

| **Measured and Ratiometric Parameters by Strain** |  |  |  |  |  |  |  |
| --- | --- | --- | --- | --- | --- | --- | --- |
| **Enzyme rates (pmol/min/mg tissue)** |  |  |  |  |  |  |  |
| Basal TKT | Mean±SE | Tukey group | N | Supplemented TKT | Mean±SE | Tukey group | n |
| ASN | 375.6±29.9 | A | 6 | ASN | 2412.1±340.3 | A | 6 |
| LT | 352.1±43.4 | A | 6 | LT | 3324.2±515.0 | A | 6 |
| TEM | 203.2±51.8 | B | 6 | TEM | 1744.5±511.3 | A | 6 |
| TXD | 363.1±10.2 | A | 6 | TXD | 3270.1±338.1 | A | 6 |
| HRN | 359.2±27.7 | A | 6 | HRN | 2928.6±238.2 | A | 6 |
| One-way ANOVA *p* |  | **0.0113\*** |  | One-way ANOVA *p* |  | 0.0537 |  |
| G6PDH | Mean±SE | Tukey group | n | GR | Mean±SE | Tukey group | n |
| ASN | 12529.7±1535.7 | B | 6 | ASN | 1593.8±111.7 | ABC | 6 |
| LT | 18589.9±1806.7 | AB | 6 | LT | 1189.8±80.0 | C | 6 |
| TEM | 22129.8±2446.4 | A | 6 | TEM | 1412.3±87.1 | BC | 6 |
| TXD | 18152.9±1293.1 | AB | 6 | TXD | 2050.4±124.1 | A | 6 |
| HRN | 19604.2±1423.8 | AB | 6 | HRN | 1766.1±227.0 | AB | 6 |
| One-way ANOVA *p* |  | **0.0114\*** |  | One-way ANOVA *p* |  | **0.0019\*** |  |
| **Metabolite pools** |  |  |  |  |  |  |  |
| NADP(H) (μM) | Mean±SE | ANOVA | n | GSH+GSSG (mM) | Mean±SE | Tukey group | n |
| ASN | 8.23±1.32 | B | 5 | ASN | 1.35±0.04 | A | 6 |
| LT | 9.35±0.95 | B | 6 | LT | 1.33±0.03 | A | 6 |
| TEM | 6.34±0.34 | B | 6 | TEM | 0.73±0.07 | C | 6 |
| TXD | 12.24±1.43 | AB | 5 | TXD | 1.42±0.05 | A | 6 |
| HRN | 17.02±2.97 | A | 6 | HRN | 0.99±0.06 | B | 6 |
| One-way ANOVA *p* |  | **0.0014\*** |  | One-way ANOVA *p* |  | **<0.0001\*** |  |
| **Morphometric parameters** |  |  |  |  |  |  |  |
| [Protein] (mg/mL) | Mean±SE | Tukey group | n | Liver mass (mg) | Mean±SE | Tukey group | n |
| ASN | 17.92±1.32 | B | 6 | ASN | 300.0±47.6 | A | 6 |
| LT | 26.94±0.87 | A | 6 | LT | 253.9±56.3 | A | 6 |
| TEM | 23.61±0.92 | A | 6 | TEM | 255.7±64.2 | A | 5 |
| TXD | 25.28±1.76 | A | 6 | TXD | 434.6±12.5 | A | 6 |
| HRN | 18.27±0.61 | B | 6 | HRN | 419.6±94.1 | A | 6 |
| One-way ANOVA *p* |  | **<0.0001\*** |  | One-way ANOVA *p* |  | 0.1003 |  |
| HSI (%) | Mean±SE | Tukey group | n | Length (mm) | Mean±SE | Tukey group | n |
| ASN | 0.910±0.073 | A | 6 | ASN | 154.8±8.5 | AB | 6 |
| LT | 0.752±0.066 | A | 6 | LT | 153.3±7.3 | AB | 6 |
| TEM | 0.901±0.125 | A | 5 | TEM | 145.7±10.6 | B | 6 |
| TXD | 0.860±0.037 | A | 6 | TXD | 176.0±2.4 | A | 6 |
| HRN | 1.075±0.187 | A | 6 | HRN | 158.3±4.4 | AB | 6 |
| One-way ANOVA *p* |  | 0.3951 |  | One-way ANOVA *p* |  | 0.0780 |  |
| Weight (g) | Mean±SE | Tukey group | n |  |  |  |  |
| ASN | 33.33±4.64 | AB | 6 |  |  |  |  |
| LT | 33.00±5.60 | AB | 6 |  |  |  |  |
| TEM | 28.67±6.43 | B | 6 |  |  |  |  |
| TXD | 50.83±1.96 | A | 6 |  |  |  |  |
| HRN | 38.17±3.42 | AB | 6 |  |  |  |  |
| One-way ANOVA *p* |  | **0.0260\*** |  |  |  |  |  |
| **Ratiometric relationships** |  |  |  |  |  |  |  |

| TKT Latency (%) | Mean±SE | Tukey group | n | (GSH+GSSG)/GR<br>(pmol/pmol/min) | Mean±SE | Tukey group | n |
| --- | --- | --- | --- | --- | --- | --- | --- |
| ASN | 83.2±2.2 | A | 6 | ASN | 2.61±0.23 | AB | 6 |
| LT | 88.7±1.2 | A | 6 | LT | 3.44±0.26 | A | 6 |
| TEM | 87.5±2.1 | A | 6 | TEM | 1.62±0.23 | C | 6 |
| TXD | 88.4±1.1 | A | 6 | TXD | 2.12±0.15 | BC | 6 |
| HRN | 87.6±0.6 | A | 6 | HRN | 1.83±0.24 | BC | 6 |
| One-way ANOVA *p* |  | 0.1203 |  | One-way ANOVA *p* |  | **<0.0001\*** |  |
| G6PDH/basal TKT | Mean±SE | Tukey group | n | G6PDH/<br>supplemented TKT | Mean±SE | Tukey group | n |
| ASN | 34.47±4.83 | B | 6 | ASN | 5.58±0.77 | B | 6 |
| LT | 54.67±4.49 | B | 6 | LT | 6.08±0.70 | B | 6 |
| TEM | 148.74±42.06 | A | 6 | TEM | 16.71±3.92 | A | 6 |
| TXD | 50.31±4.11 | B | 6 | TXD | 5.73±0.50 | B | 6 |
| HRN | 57.60±9.23 | B | 6 | HRN | 7.04±1.09 | B | 6 |
| One-way ANOVA *p* |  | **0.0029\*** |  | One-way ANOVA *p* |  | **0.0011\*** |  |
| Liver soluble<br>protein/liver mass<br>(µg protein/g liver) | Mean±SE | Tukey group | n | Liver soluble<br>protein/body mass<br>(µg protein/g fish) | Mean±SE | Tukey group | n |
| ASN | 53.77±3.95 | B | 6 | ASN | 482.7±41.2 | A | 6 |
| LT | 80.82±2.60 | A | 6 | LT | 605.9±51.8 | A | 6 |
| TEM | 70.81±2.75 | A | 6 | TEM | 643.4±84.4 | A | 6 |
| TXD | 75.84±5.28 | A | 6 | TXD | 649.2±44.6 | A | 6 |
| HRN | 54.81±1.82 | B | 6 | HRN | 588.2±105.2 | A | 6 |
| One-way ANOVA *p* |  | **<0.0001\*** |  | One-way ANOVA *p* |  | 0.451 |  |

Analyses were conducted using one-way ANOVAs, followed by Tukey's multiple comparisons tests. Means not sharing a letter differ significantly by Tukey's multiple comparisons test ( $p < 0.05$ ).

Body mass differed among strains (one-way ANOVA, *p=*0.026), with the TXD strain exhibiting significantly greater mass than the TEM strain, whereas the other strains showed intermediate values. Similar trends were observed in body length, although the difference was not statistically significant. Consistent with prior work using fish reared in the same tanks, the ASN and TEM strains amassed less weight for a given length, relative to LT and HRN, with the greatest observed for TXD.[39] The liver mass was substantially lower for the LT strain (254±57 mg) than for the TXD strain (435±13 mg), though the difference was not statistically significant, as other strains yielded intermediate values. The HSI did not differ significantly between strains but trended lowest for LT and highest for the TEM strain.

The mean soluble protein concentration across all strains ranged from 17.9 to 26.9 mg/mL, consistent with those determined in lake trout (16.6-19.8 mg/mL).[33] The extractable protein concentration differed significantly between strains (one-way ANOVA, p<0.0001), with ASN and HRN strains exhibiting lower extractable protein than the remaining strains.

Total glutathione concentrations in liver extracts ranged from 0.72 to 1.42 mM, consistent with the expected cytosolic glutathione concentration range observed in mammals (0.5-10 mM).[40, 41] Our measurements assessed the total glutathione pool, not the reduced-to-oxidized ratio, which typically ranges from 30 to >100:1 and drops considerably under oxidative stress.[17, 42] Total glutathione pool size differed significantly among strains (one-way ANOVA, p<0.0001), with ASN, LT, and TXD strains exhibiting comparable and higher concentrations than HRN, which in turn exceeded the TEM strain (Fig. 2a). Glutathione synthesis is an ATP-dependent two-enzyme process, rate-limited by glutamate-cysteine ligase and the availability of cysteine.[23, 24, 43] Irrespective of the underlying biochemical control, the ASN, LT, and TXD strains had a significantly greater pool of glutathione available to buffer oxidative stress per unit liver tissue mass than the HRN and TEM strains.

**Fig. 2.**
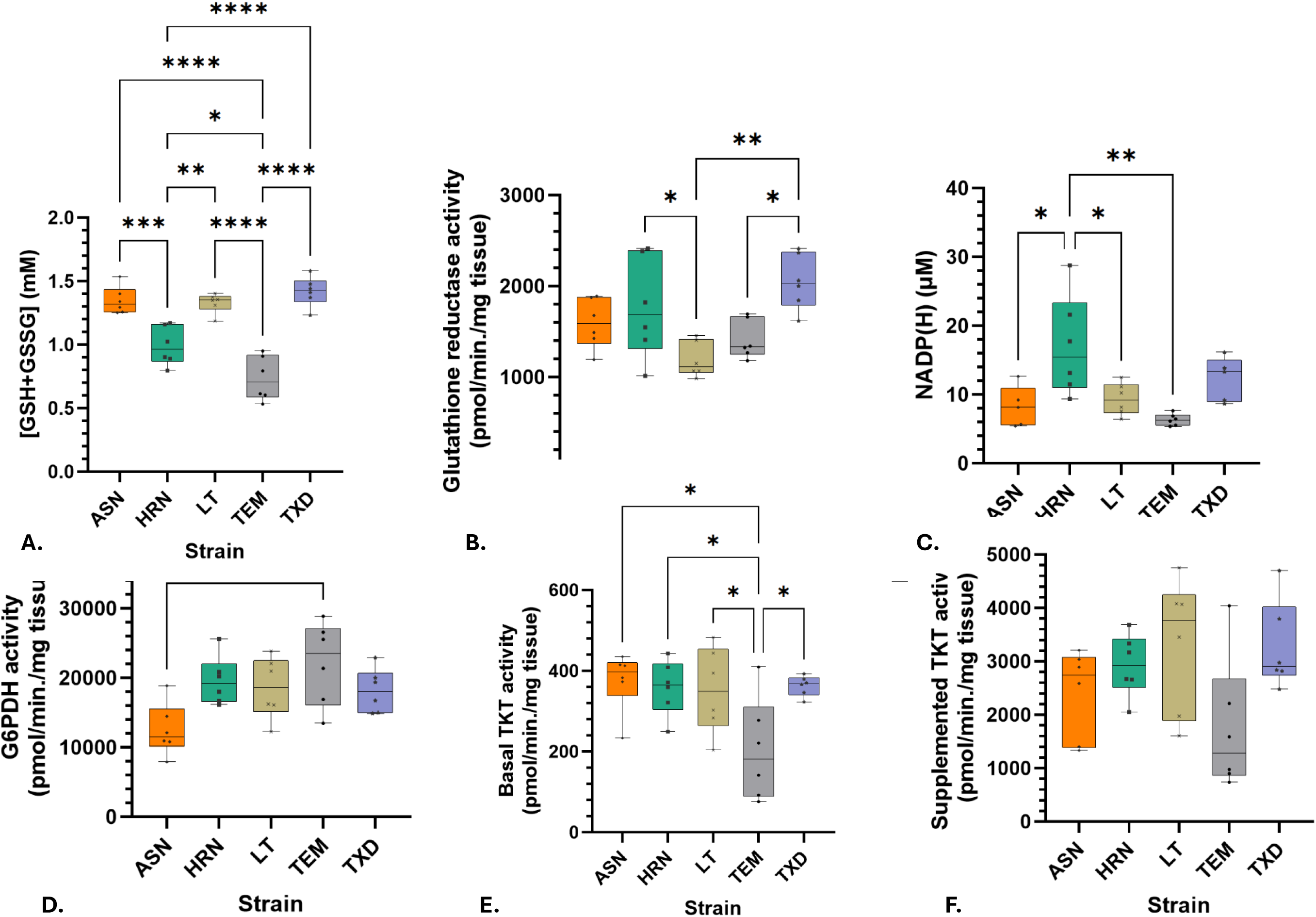
A.) total glutathione concentrations, B.) glutathione reductase activity, C.) total NADP(H) concentrations, D.) G6PDH activity, E.) Basal TKT activity, and F.) Supplemented TKT activity. Each point represents an individual fish, with n=6 per strain, except for NADP(H) for which n=5 for the TXD and ASN strains and each assay run in duplicate or triplicate per sample. Significance was tested using one-way ANOVAs with Tukey’s post hoc tests for multiple comparisons for all parameters.

As maintenance of a reduced glutathione pool depends on glutathione reductase (GR) activity, we quantified GR as an index of glutathione recycling capacity. GR activity ranged from 1,190 to 2,050 pmol/min/mg tissue, consistent with hepatic levels in hatchery-reared rainbow trout.[22] GR activity differed among strains (one-way ANOVA, *p=*0.0019), with the LT and TEM strains exhibiting significantly lower activity than the TXD strain on a mass basis (Fig. 2b). The gene expression of GR is upregulated by oxidative stress, while after translation, the formation of its active homodimer form and enzyme activity is dependent on the availability of its riboflavin-dependent cofactor flavin adenine dinucleotide (FAD), GSH-GSSG ratio, and NADPH.[17, 42] In our assays, the substrates were not limiting, and strains consumed the same diets, suggesting that differences in activity reflect differences in GR expression, affinity, or stability. The TXD strain had a significantly greater GR capacity per unit liver tissue mass, and thus a greater ability to recycle available glutathione to its reduced form, than the LT or TEM strain.

As GR activity is dependent on NADPH availability in vivo, we quantified total NADP(H) concentrations as an index of the supply of reducing equivalents. We note that the measured NADP(H) represents the total pool, which rapidly interconverts between oxidized and reduced forms; therefore, our measurements do not directly reflect reducing power. Total NADP(H) concentrations ranged from 6.3 μM to 17.0 μM across strains. At ambient temperature, NADP(H) concentrations differed significantly amongst strains (one-way ANOVA, *p=*0.0014), with the TEM strain exhibiting the lowest concentration (6.3 μM) which, together with LT and ASN strains, was significantly lower than that of the HRN strain (17.0 μM) (Fig. 2c). The HRN strain had a significantly larger NADP(H) pool available, and thus a greater potential for NADPH-dependent biosynthesis or antioxidant defense per unit liver tissue mass.

To evaluate whether observed differences in NADP(H) availability were supported by upstream capacity for reducing-equivalent generation, we quantified G6PDH activity as an index of oxidative PPP capacity. NADPH can be generated from NADP^+^ by multiple pathways, including the oxidative PPP (G6PDH and 6PGDH) and auxiliary cytosolic enzymes such as malic enzyme and NADP-dependent isocitrate dehydrogenase.[44] G6PDH is the rate-limiting step for the oxidative PPP, and hence was used here as a pathway-specific indicator of oxidative PPP capacity. Mean hepatic G6PDH activity ranged from 12,530 to 22,130 pmol/min/mg tissue, consistent with reported hepatic G6PDH activities in rainbow trout.[44] G6PDH activities differed among strains (one-way ANOVA, *p=*0.0114), with the ASN strain exhibiting significantly lower activity than the TEM strain. Other strains exhibited intermediate values (Fig. 2d). In addition to transcriptional regulation, post-translational modifications affect the expression, stability, and activity of G6PDH.[45] G6PDH also undergoes a concentration- and NADPH-dependent conversion from active dimers and tetramers to inactive monomers, with NADPH being a potent inhibitor.[45–47] Irrespective of the underlying biochemical control, the TEM strain had a significantly greater G6PDH capacity on a functional basis and, thus, a greater maximal oxidative PPP capacity than the ASN strain per mass of liver tissue.

As oxidative PPP governs NADPH supply but does not determine downstream carbon redistribution, we next evaluated TKT activity to assess strain-specific differences in non-oxidative PPP capacity. Mean basal TKT activity ranged from 203 to 376 pmol/min/mg tissue, near the upper end of values reported for lake trout reared at 9°C.[33] Basal TKT activity differed among strains (one-way ANOVA, *p=*0.0113), with the TEM strain exhibiting significantly lower activity than all other strains, which were otherwise comparable on a tissue-normalized basis (Fig. 2e). Thiamine diphosphate (TDP)-supplemented TKT activities ranged from 1,744 to 3,324 pmol/min/mg tissue and similarly exceeded reported values for lake trout,[33] though were more variable between individuals (Fig. 2f). Although the TEM strain continued to exhibit the lowest activity amongst strains, this did not reach statistical significance (*p=*0.0537). TKT latency, reflecting the proportion of inactive enzymes under basal conditions, did not differ significantly among strains. Thiamine levels have been reported to affect TKT expression,[48] and post-translational modifications affect its activity.[49] TKT further undergoes a concentration-, TDP-, and Mg^2+^-dependent conversion from inactive monomers to active dimers.[33, 50] Irrespective of the underlying biochemical control, the TEM strain had a significantly lower TKT capacity on a functional basis, and thus a more limited ability per mass of liver tissue to use the non-oxidative PPP than other strains.

### Strain-specific metabolic organization

Overall, the relative biochemical profiles differed markedly among strains, with each exhibiting a distinct pattern of metabolic organization. To visualize the collective biochemical phenotype of each strain, measured parameters were normalized as z-scores and summarized as a heatmap (Fig. 3). The heatmap highlights coordinated differences among strains that are less apparent when parameters are considered individually. Relative to other strains, ASN exhibited the lowest hepatic soluble protein density, low TDP-supplemented TKT activity, one of the lowest NADP(H) pools, and the lowest oxidative PPP capacity. However, it exhibited a relatively high basal TKT activity and glutathione pool. HRN similarly had a low soluble protein density, but the largest NADP(H) pool, a comparatively small glutathione pool, and intermediate PPP activity. LT, by contrast, had the highest protein density, but had an elevated total glutathione pool on par with that of ASN, but with low GR activity. The differences between TEM and its cross (TXD) were the most striking. The non-oxidative capacity of TEM was the lowest among all strains, with the lowest NADP(H) pool, glutathione pool, and second-to-lowest GR activity. TEM, however, had the highest oxidative PPP capacity of all strains. TXD, by contrast, had the highest GR capacity, the largest glutathione pool, and intermediate capacities for both the oxidative and non-oxidative PPP.

**Fig. 3.**
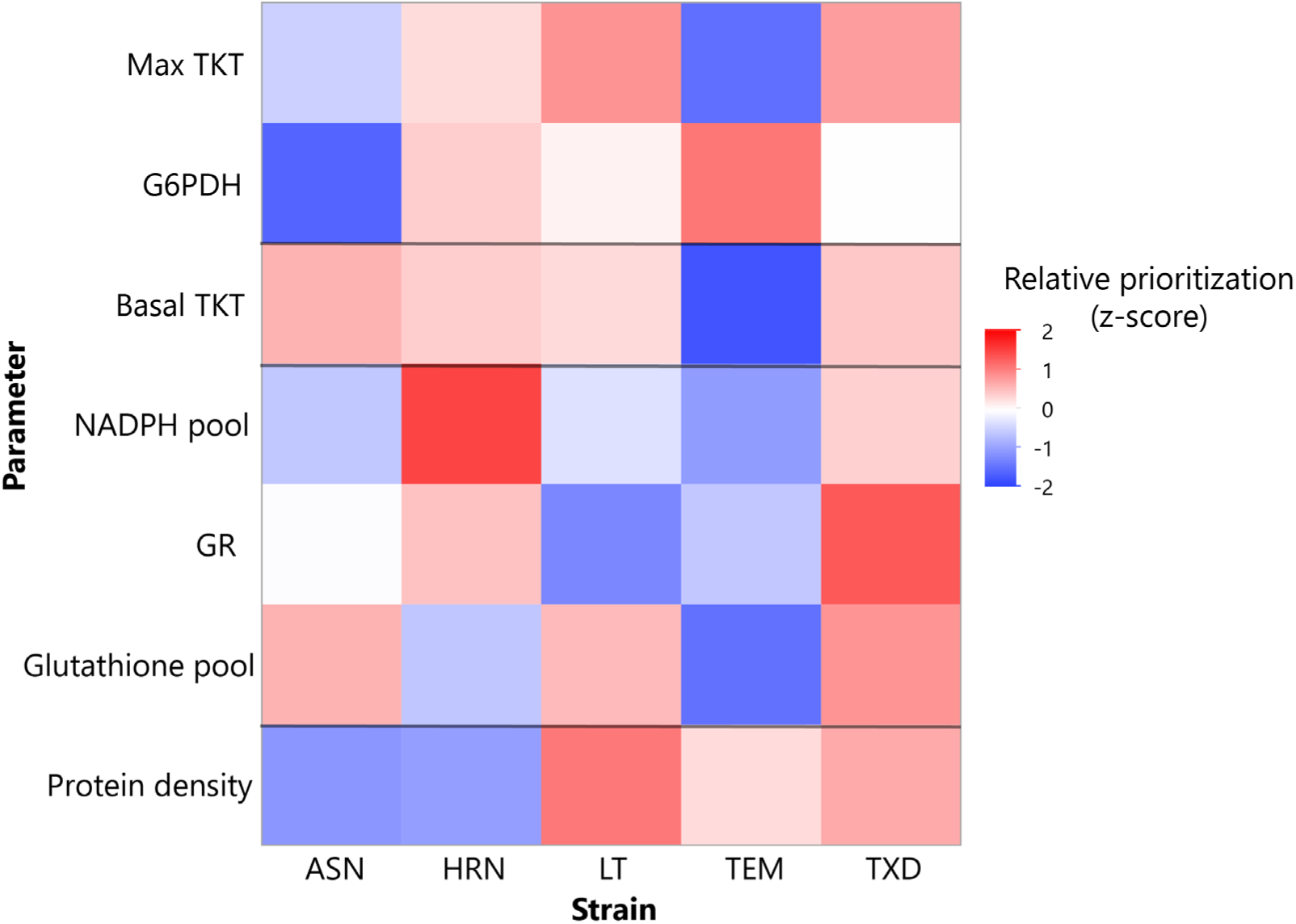
Z-scored heat map showing relative differences in tissue-normalized metabolic markers among brook trout strains. Within each measured parameter (rows), red indicates values above the mean across strains and blue indicates values below the mean. Max TKT refers to TDP-supplemented TKT activity, the NADP(H) pool represents total NADP^+^ and NADPH, and the glutathione pool represents total GSH and GSSG.

### Using ratiometric relationships to identify metabolic organization

Given the complexity of both the PPP and glutathione axes, we condensed the data into ratios that reflect varying levels of investment in biochemical pathways. The rationale, description, and interpretation of these ratiometric parameters are tabulated in Table 2.

**Table 2.**
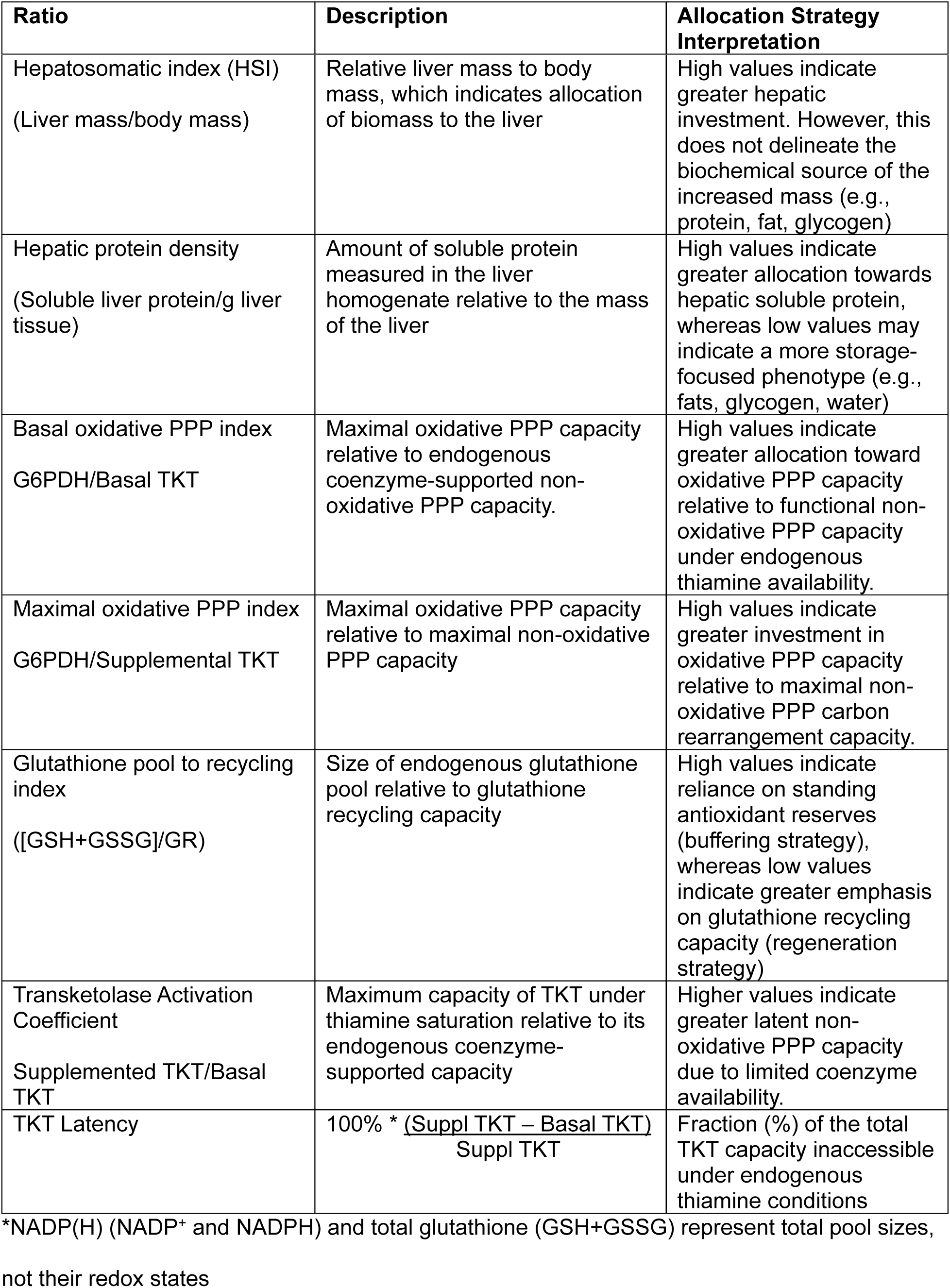
Ratiometric framework for interpreting metabolic allocation strategies.

| Ratio | Description | Allocation Strategy Interpretation |
| --- | --- | --- |
| Hepatosomatic index (HSI)<br>(Liver mass/body mass) | Relative liver mass to body mass, which indicates allocation of biomass to the liver | High values indicate greater hepatic investment. However, this does not delineate the biochemical source of the increased mass (e.g., protein, fat, glycogen) |
| Hepatic protein density<br>(Soluble liver protein/g liver tissue) | Amount of soluble protein measured in the liver homogenate relative to the mass of the liver | High values indicate greater allocation towards hepatic soluble protein, whereas low values may indicate a more storage-focused phenotype (e.g., fats, glycogen, water) |
| Basal oxidative PPP index<br>G6PDH/Basal TKT | Maximal oxidative PPP capacity relative to endogenous coenzyme-supported non-oxidative PPP capacity. | High values indicate greater allocation toward oxidative PPP capacity relative to functional non-oxidative PPP capacity under endogenous thiamine availability. |
| Maximal oxidative PPP index<br>G6PDH/Supplemental TKT | Maximal oxidative PPP capacity relative to maximal non-oxidative PPP capacity | High values indicate greater investment in oxidative PPP capacity relative to maximal non-oxidative PPP carbon rearrangement capacity. |
| Glutathione pool to recycling index<br>([GSH+GSSG]/GR) | Size of endogenous glutathione pool relative to glutathione recycling capacity | High values indicate reliance on standing antioxidant reserves (buffering strategy), whereas low values indicate greater emphasis on glutathione recycling capacity (regeneration strategy) |
| Transketolase Activation Coefficient<br>Supplemented TKT/Basal TKT | Maximum capacity of TKT under thiamine saturation relative to its endogenous coenzyme-supported capacity | Higher values indicate greater latent non-oxidative PPP capacity due to limited coenzyme availability. |
| TKT Latency | $100\% * \frac{(\text{Suppl TKT} - \text{Basal TKT})}{\text{Suppl TKT}}$ | Fraction (%) of the total TKT capacity inaccessible under endogenous thiamine conditions |

\*NADP(H) (NADP<sup>+</sup> and NADPH) and total glutathione (GSH+GSSG) represent total pool sizes, not their redox states

Ratios of G6PDH to basal TKT activity were used to evaluate relative oxidative versus non-oxidative PPP capacity. In all cases, the G6PDH activity exceeded basal TKT activity. This ratio ranged from 20.7 to 336.0 across strains (n=30, mean=69.2±11.1, median=52.0). It differed significantly among strains (one-way ANOVA, *p=*0.0029), with TEM exhibiting substantially higher values than all other strains, indicating a strong bias towards oxidative PPP relative to endogenous cofactor-supported carbon rearrangement capacity. This is a function of both the composite higher G6PDH activity and the lower basal TKT activity in TEM, as shown in Fig. 2. When G6PDH was normalized to TDP-supplemented TKT activity (termed the maximal oxidative PPP index), strain differences were retained (*p=*0.0011), with TEM again exhibiting the highest ratio. This ratio ranged from 3.37 to 34.50 across strains (n=30, mean=8.23±1.12, median=6.49).

Finally, ratios of glutathione pool size to GR activity (termed the glutathione pool to recycling index) were used to assess the constraints on redox buffering and recycling. The (GSH+GSSG)/GR ratio ranged from 0.96 to 4.13 across strains (n=30, mean=2.32±0.15, median=2.27). This ratio differed significantly among strains (one-way ANOVA, p<0.0001), with LT exhibiting the highest ratio, TEM the lowest, and ASN and TXD intermediate ratios. For the LT strain, this is a function of both its lower GR activity and relatively high total glutathione concentrations, shown in Fig. 2.

### Evaluating the effect of body size on measured parameters

After accounting for strain, neither body length nor body mass significantly explained variation in any measured or derived ratiometric biochemical index except G6PDH (Table 3.) Conversely, liver mass variation was not significantly explained by strain, but was by body size. The positive association of G6PDH activity with body size suggests that oxidative PPP capacity may continue to scale with somatic growth, even though most hepatic metabolic traits appear largely independent of body size within the size range examined.

**Table 3.** Standard least squares models evaluating the effects of strain and body size on biochemical parameters.

|  | Strain versus length |  | Strain versus mass |  |
| --- | --- | --- | --- | --- |
|  | Strain (p) | Length (p) | Strain (p) | Mass (p) |
| Liver mass | 0.3434 | **0.0041** | 0.5280 | **0.0021** |
| Protein density | **<0.0001** | 0.6377 | **<0.0001** | 0.5040 |
| Basal TKT | **0.018** | 0.775 | **0.020** | 0.891 |
| TDP-supplemented TKT | 0.131 | 0.433 | 0.133 | 0.531 |
| G6PDH | **0.0001** | **0.0002** | **0.0002** | **0.0005** |
| NADP(H) | **0.0017** | 0.4625 | **0.0011** | 0.2440 |
| GR | **0.0124** | 0.5161 | **0.0187** | 0.5324 |
| Total glutathione | **<0.0001** | 0.2105 | **<0.0001** | 0.2811 |
| (GSH+GSSG)/GR | **<0.0001** | 0.7823 | **<0.0001** | 0.7823 |
| G6PDH/basal TKT | **0.0007** | 0.0594 | **0.0008** | 0.0689 |
| G6PDH/supp TKT | **0.0010** | 0.2888 | **0.0009** | 0.2672 |

## Discussion

Although brook trout thermal physiology, reproductive biology, and ecology have been comprehensively examined,[51] little is known about strain-specific differences in the biochemical organization of central carbon metabolism and redox homeostasis. Instead, studies have focused on understanding differences in the catchability, growth and survival of strains in the wild.[52],[53, 54] Our analysis uses biochemical metrics associated with PPP capacity and glutathione metabolism that reveal differences in baseline metabolic allocation strategies among brook trout strains. By examining fish reared under identical ambient environmental conditions, we identify contrasting metabolic phenotypes that link organismal and molecular observations to the underlying biochemical architecture of each strain. These metabolic phenotypes provide a foundation to understand strain-specific responses to two key contemporary environmental stressors: warming and hypoxia. This approach can also be applied to diverse organisms in which the PPP and glutathione-dependent metabolism are conserved, plus provides a conceptual framework to understand recycling strategies in organisms using other low-molecular-weight thiol systems.

### Hepatic soluble protein density versus hepatosomatic index

Liver mass increased with body size (length, *p=*0.004, mass, *p=*0.002) but did not differ among strains after accounting for body size (length model, *p=*0.343 and weight model, *p=*0.528). Conversely, hepatic soluble protein density differed significantly among the strains (p<0.0001) but was independent of body length (*p=*0.638) or weight (*p=*0.504), indicating that strain differences in protein density reflected hepatic composition rather than fish size.

Despite exhibiting comparable HSI values, the ASN and HRN strains showed substantially lower hepatic soluble protein densities than other strains, indicating reduced soluble protein density rather than reduced liver investment (Fig. 4). Conversely, LT exhibited a higher soluble protein concentration despite a relatively low HSI. As approximately 50% of the proteins in hepatocytes are enzymes,[55] the high protein concentration and low HSI of LT is consistent with a protein-dense hepatic phenotype in which metabolic machinery is concentrated rather than expanded through increased organ mass. TXD and TEM occupied intermediate positions, although TXD showed greater variability in protein density (Fig. 4).

**Fig. 4.**
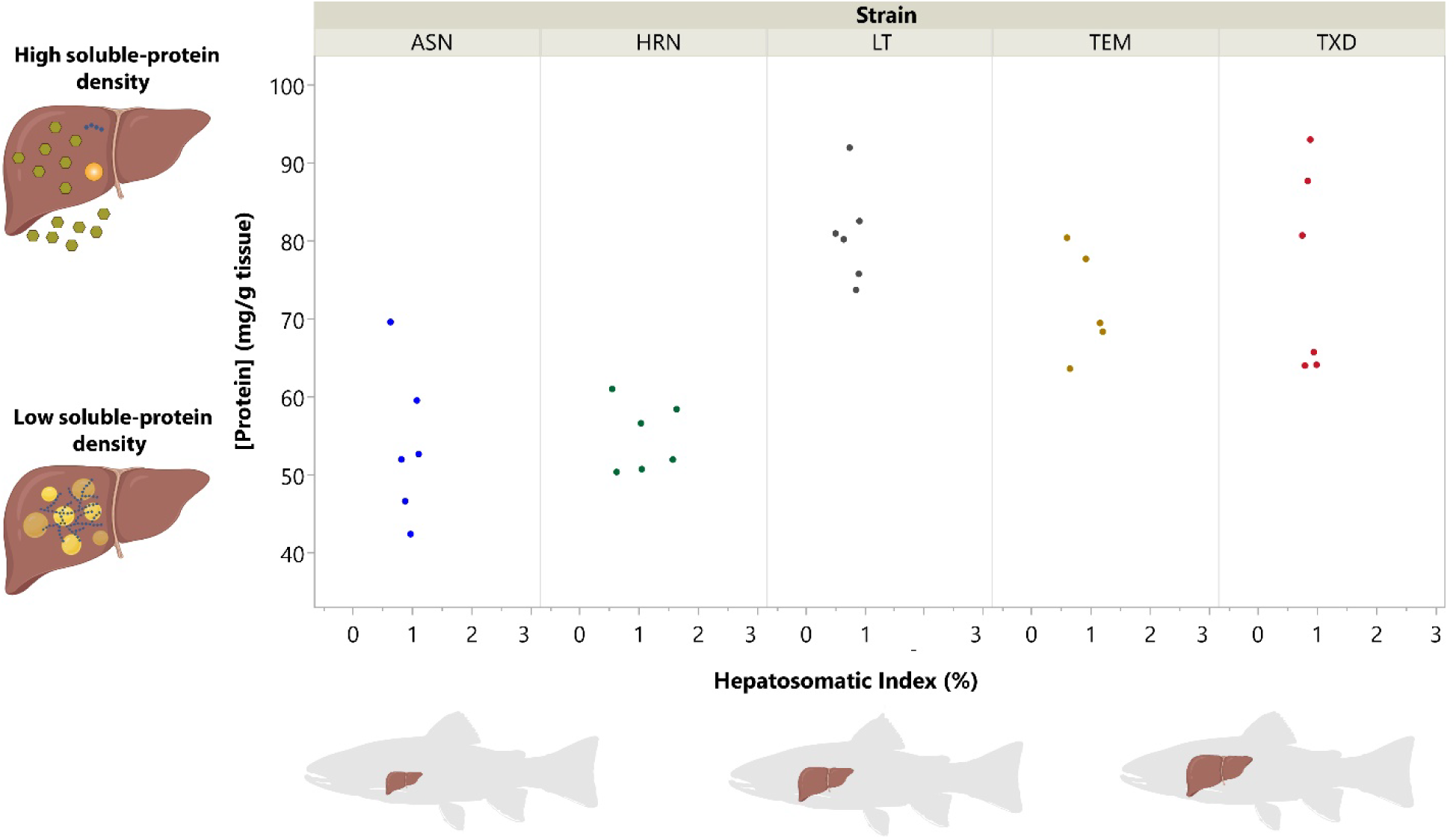
Relationship between soluble hepatic protein density (mg/g liver tissue) and hepatosomatic index (liver mass/body mass (%)) across strains. Each point represents an individual fish.

Soluble protein density was significantly higher in the TEM, TXD, and LT strains (71-81 mg protein/g liver) than in the HRN and ASN strains (54-55 mg protein/g liver) (Table 2). Given the lack of size dependence and since all strains were reared under identical conditions, differences in soluble protein concentration likely reflect inherent differences in hepatic composition rather than environmental effects. The mild protein extraction conditions (e.g., extraction into isosmotic sucrose and 12,000 × g centrifugation) would be expected to enrich the supernatant in soluble cytosolic components, while depleting insoluble structural proteins, intact organelles, nuclei, and large membrane fragments. Consequently, our measurements do not represent total liver protein, often cited as ∼16-20% of liver mass, but rather a soluble fraction enriched in cytosolic proteins associated with intermediary metabolism.

Higher soluble protein concentrations suggest a greater proportion of the liver biomass is devoted to soluble proteins, including metabolic enzymes, whereas lower densities are consistent with a larger contribution from non-protein components (e.g., glycogen, lipids, water), possibly suggesting a greater emphasis on storage. However, the underlying physiological basis for these differences remains unresolved and would benefit from future measurements of hepatic lipid, glycogen, water content, and proteomic composition.

### Relative levels of protein, glutathione, NADPH, and enzyme activity by strain

Divergent metabolic strategies among brook trout strains were evident under common ambient conditions that could influence observed differences in growth, feeding behavior, and catchability of these same brook trout strains reared in a common garden experiment.[39] The strains separate broadly along three recurring metabolic themes: antioxidant buffering, oxidative PPP investment, and deployable non-oxidative PPP capacity (Fig. 3). ASN exhibited a buffering-oriented phenotype, characterized by elevated glutathione pools and higher basal TKT activity, but relatively low oxidative and maximal non-oxidative PPP capacity, NADP(H) availability, and protein density. This pattern suggests a metabolic strategy that emphasizes antioxidant buffering and baseline non-oxidative PPP activity rather than high regenerative capacity or maximal pathway capacity. The LT strain displayed a protein-dense hepatic phenotype with larger glutathione pools relative to recycling capacity, similarly consistent with a strategy emphasizing redox buffering. The TEM strain, by contrast, displayed the highest oxidative PPP capacity but limited non-oxidative PPP deployment and capacity, along with relatively low NADP(H) levels, glutathione pool size, and glutathione recycling capacity. This profile indicates a strategy biased toward NADPH generation through G6PDH, with limited investment in non-oxidative PPP or antioxidant buffering. The HRN strain maintained the largest NADP(H) pool and exhibited moderate capacities for oxidative and non-oxidative PPP activity. Its relatively limited glutathione reserves, combined with its moderate glutathione recycling capacity, are consistent with a turnover-oriented redox strategy that favors regeneration of reducing equivalents over antioxidant storage. The TXD strain exhibited a phenotype consistent with greater investment in oxidative-stress defense and detoxification, reflected by its elevated GR capacity and large glutathione pool. It maintains moderate basal TKT activity and NADPH pool availability, along with high TKT capacity but only intermediate oxidative PPP capacity. Together, this profile suggests a strategy oriented toward downstream detoxification and redox resilience rather than for maximizing carbon flux through the PPP. Collectively, these patterns indicate that strains differ not only in metabolic competence but also in how liver resources are allocated among buffering, regeneration, and capacity, potentially reflecting adaptation to distinct ecological and physiological demands.

### Ratiometric indices

Although strain differences were apparent across enzyme activities and redox pools, these comparisons alone do not resolve how strains differentially handle glutathione recycling or employ the PPP. For example, an elevated total glutathione level in isolation does not speak to the ability of the organism to recycle glutathione to its active reduced form. Since PPP function reflects coordinated investment in reserve capacity, oxidative flux, and redox buffering, we integrated these dimensions to define strain-specific metabolic strategies. These were structured around the emphasis on:

- glutathione pool generation versus its recycling capacity (GSH+GSSG)/GR),
- the relative emphasis of the oxidative PPP versus the non-oxidative PPP (G6PDH/TKT), and
- the total TKT capacity available if TDP were unlimiting versus the capacity accessible under endogenous TDP cofactor availability (TDP-supplemented TKT/basal TKT).

Ratiometric activity indices, such as supplemented TKT-to-basal TKT,[56] G6PDH-to-pyruvate kinase,^44^ or G6PDH-to-6-phosphogluconate dehydrogenase,^45^ can provide complementary, or in some cases improved, diagnostic information relative to absolute activities. While various presentations of TDP-supplemented/basal TKT activity are commonly employed, neither the glutathione pool-to-glutathione recycling capacity nor the G6PDH-to-TKT activity ratios have been reported previously. By integrating oxidative capacity (G6PDH), non-oxidative carbon rearrangement (TKT), and glutathione buffering relative to recycling capacity, we identify distinct patterns of hepatic metabolic organization. These patterns do not reflect differences in realized flux per se, but rather differences in pathway capacity and redox infrastructure that may influence how strains respond to oxidative challenge, nutrient variability, or growth demands.

Maintenance of a glutathione pool requires dietary amino acid uptake or endogenous synthesis and sequestration, as well as ATP for glutathione formation,[57],[58] representing a quantifiable metabolic allocation. In environments where oxidative stress or xenobiotic exposure is limited, such allocation toward glutathione buffering depletes resources that could be better spent on other uses and may be reduced. Differences in glutathione pool size relative to recycling capacity, therefore, reflect not only redox demand but also broader allocation strategies within hepatic metabolism.

For all strains, the total glutathione pool-to-GR ratio exceeded one, with HRN and TEM having the lowest ratios, followed by TXD, then ASN, and LT exhibiting the highest ratio (Fig. 5). Prior studies have associated higher total glutathione[59] and higher GR activity with greater pollution exposure[60] and higher dietary lipids,[22] with seasonal and gender differences contributing to observed variation.[59] Given that all fish were sampled at the same time and had experienced the same environment, the larger relative glutathione pool size in the LT strain may indicate a greater standing buffering reserve available to counter oxidative stress.

**Fig. 5.**
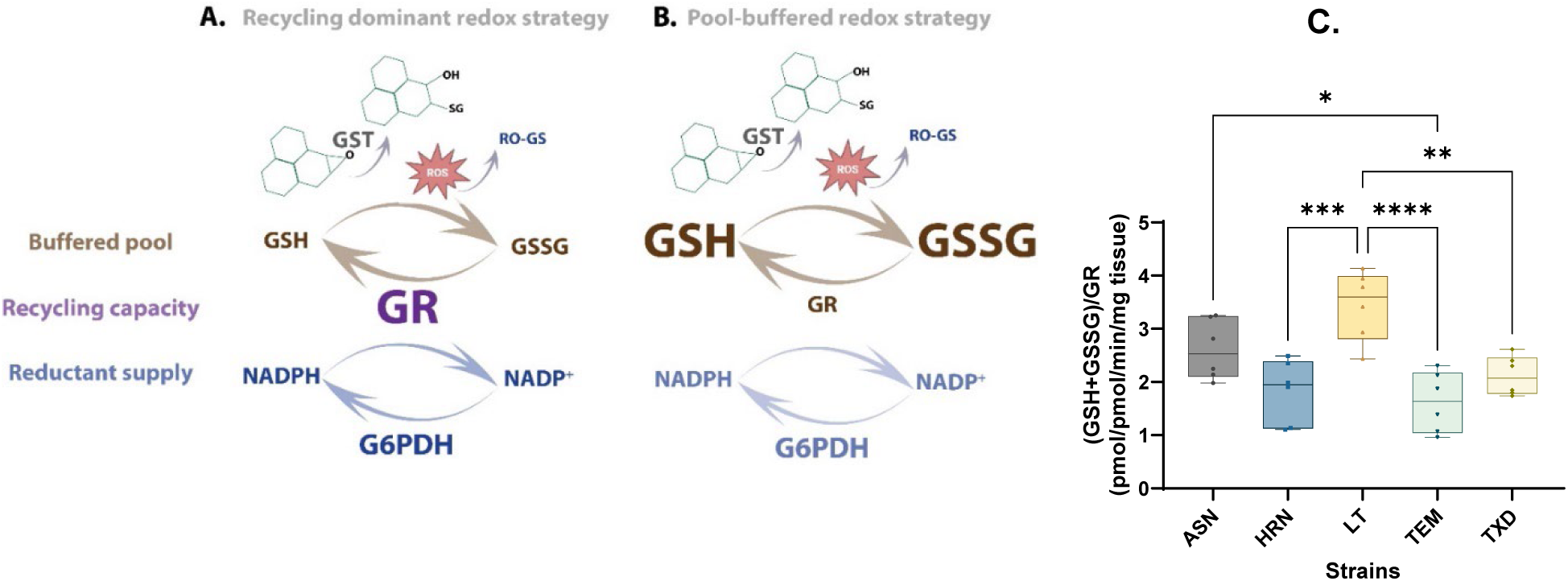
Alternative glutathione buffering versus recycling strategies underlying strain-specific metabolic phenotypes. A) glutathione recycling strategy with smaller total glutathione pools (GSH_(red)_ and GSSG_(ox)_) and heavier reliance on glutathione reductase (GR). B) glutathione buffering strategy with larger glutathione pools and reduced reliance on recycling, and C.) Ratio of (GSH+GSSG)/GR activity by strain. The box-and-whisker plots represent the median, minimum, and maximum, with each data point representing one fish, with values calculated from the average of duplicate or triplicate measurements for each assay. Strain differences were compared by one-way ANOVA followed by Tukey’s multiple comparison test (* p<0.05, ** p<0.01, *** p<0.001, ****p<0.0001).

The relative investment in the PPP and its composite arms is of interest in tumor proliferation, where upregulation of the PPP can limit cancer cells’ exposure to oxidative stress while supplying ribose and reducing equivalents to support proliferative growth.[61] Silencing G6PDH or TKT has been shown to alter ROS production and fatty acid synthesis, highlighting how PPP partitioning influences redox balance, biosynthetic capacity, and growth strategy.[62] Thus, understanding this balance is important for maintaining environmental resilience and growth characteristics in fish. G6PDH supports NADPH production for biosynthesis and antioxidant defense, suggesting its capacity is influenced by growth-related metabolic demands, though the development and regulation of hepatic G6PDH in fish remains poorly understood and warrants further investigation.

For all strains, G6PDH activity exceeded basal TKT activity, yielding G6PDH/basal TKT ratios greater than one (Fig. 6). This indicates greater relative enzymatic capacity for oxidative than endogenous cofactor-supported non-oxidative PPP capacity, favoring NADPH-generating capacity while incurring the carbon loss associated with CO_2_ release. However, the TEM strain placed significantly greater emphasis on the oxidative PPP compared to the non-oxidative phase than other strains, most notably the ASN strain. The TEM phenotype is consistent with greater investment in oxidative PPP capacity despite relatively small standing NADP(H) and glutathione pools. This pattern may reflect preferential routing of NADPH toward lipid remodeling, xenobiotic metabolism, or other NADPH-consuming anabolic processes rather than an expansion of standing antioxidant reserves. In contrast, ASN appears to allocate more resources to constitutive glutathione buffering and to endogenous cofactor-supported non-oxidative PPP capacity.

**Fig. 6.**
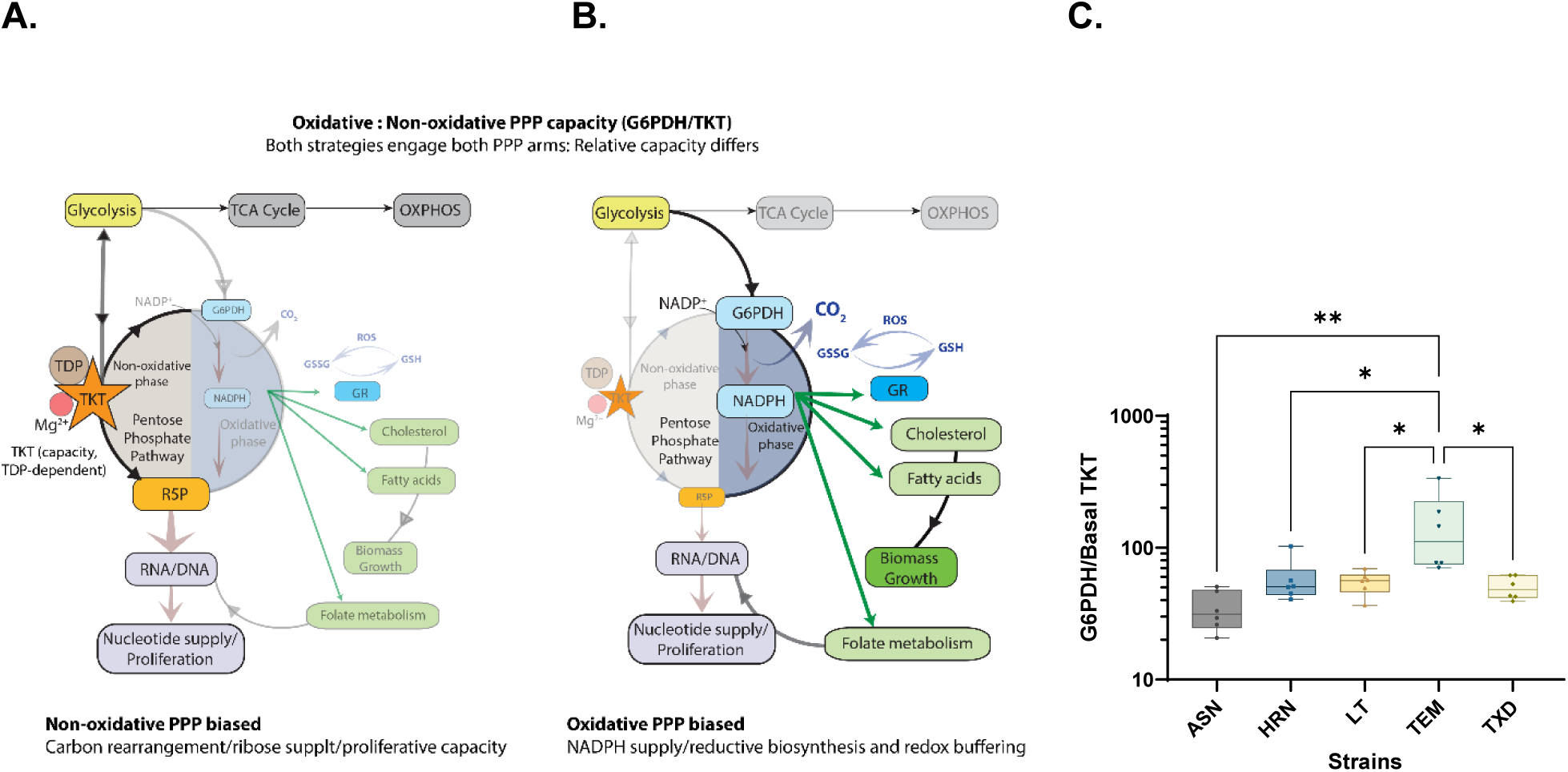
Oxidative versus non-oxidative PPP emphasis underlying strain-specific metabolic phenotypes. A.) Heavier emphasis on the non-oxidative PPP, supporting carbon flexibility and nucleotide growth capacity, B.) Heavier emphasis on the oxidative PPP, supporting NADPH supply for lipid/cholesterol synthesis, redox homeostasis, folate metabolism, and detoxification, and C.) Ratio of G6PDH/Basal TKT activity by strain. The box-and-whisker plots show the median, minimum, and maximum, with each data point representing the average of duplicate or triplicate measurements for both the G6PDH and TKT assays. Strain differences were compared by one-way ANOVA followed by Tukey’s multiple comparison test (* p<0.05, ** p<0.01).

Comparison of basal and TDP-supplemented capacities distinguishes realized enzyme capacity under endogenous TDP availability from latent catalytic capacity that becomes accessible if the TDP-coenzyme limitation was removed. Basal TKT activity represents non-oxidative PPP capacity supported by endogenous TDP, whereas TDP-supplemented TKT activity represents the maximal catalytic capacity of the tissue enzyme pool. Consequently, the ratio of supplemented to basal activities reflects the proportion of total catalytic capacity that is inaccessible under endogenous TDP availability, rather than instantaneous flux, as the assays are carried out with a substrate (R5P) excess. Differences among strains in this ratio would indicate alternative metabolic strategies in which non-oxidative PPP capacity is either constitutively supported or maintained as a latent reserve capacity under changing nutritional or environmental demands.

This ratio trended lower for the ASN strain relative to other strains, however, the differences between strains were overall not statistically significant (Fig. 7). Although the TEM strain had significantly lower basal TKT activity than the other strains, its proportional response to TDP was similar. Thus, while strains differ in their absolute capacities, they did not differ in the proportion of catalytic capacity supported by endogenous TDP.

**Fig. 7.**
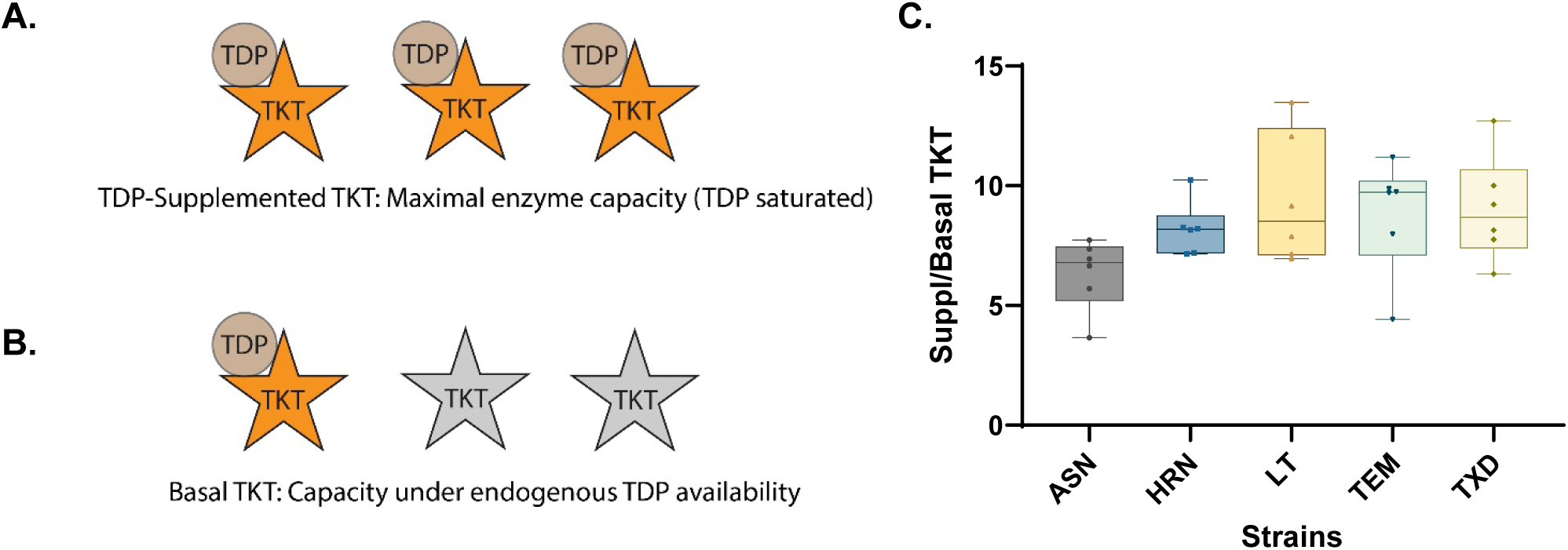
Schematic showing A.) maximal TKT activity with TKT saturated by TDP, B.) basal TKT activity under endogenous substrate-limited TDP concentrations, C.) Ratio of Supplemented TKT/Basal TKT activity by strain. The box-and-whisker plots show the median, minimum, and maximum, with each data point representing the average of triplicate measurements for the basal and supplemented TKT assays. Strain differences were compared by one-way ANOVA followed by Tukey’s multiple comparison test.

### Interpreting the network of ratiometric indices

The three-axis framework presented in Fig. 8 integrates these dimensions simultaneously, allowing strains to be positioned within a shared metabolic space defined by 1) oxidative PPP capacity relative to endogenous TDP-limited non-oxidative capacity, 2) deployable versus latent non-oxidative PPP capacity (TDP supplemented TKT/basal TKT), and 3) glutathione buffering relative to recycling potential ((GSH+GSSG)/GR) (Fig. 8.)

**Fig. 8.**
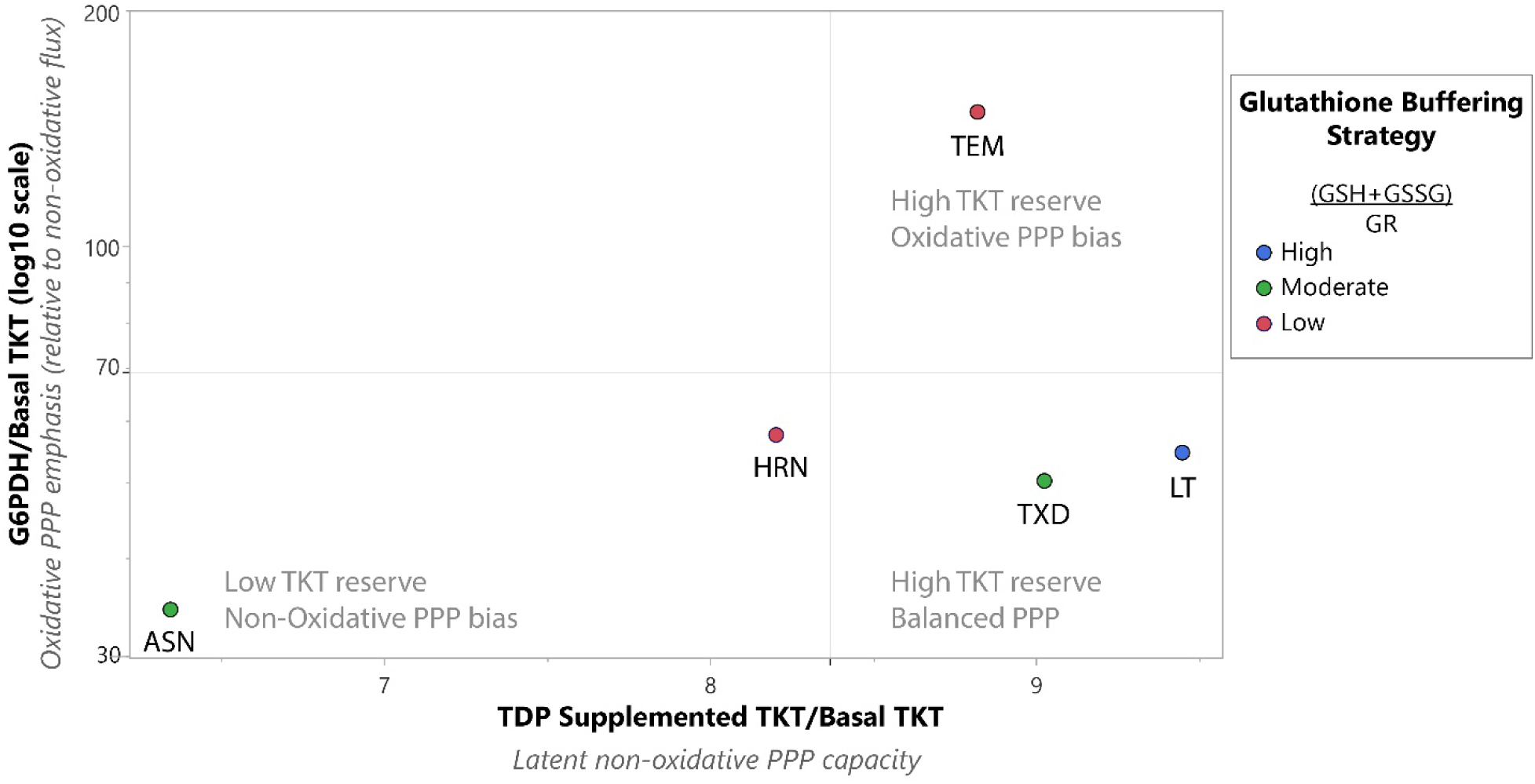
Strain-specific metabolic strategies inferred from relative pentose-phosphate pathway allocation in brook trout liver. Each point represents a brook trout strain positioned according to mean tissue-normalized ratios describing the non-oxidative and oxidative PPP capacity under ambient conditions. The x-axis (TDP-supplemented TKT/basal TKT) reflects latent non-oxidative PPP capacity relative to endogenous cofactor-supported capacity, with higher values indicating greater reserve capacity for carbon rearrangement. The y-axis (G6PDH/basal TKT (log10 scale)) reflects oxidative PPP emphasis and its NADPH-generating potential relative to endogenous cofactor-supported non-oxidative PPP capacity. Point color denotes glutathione buffering strategy based on the (GSH+GSSG)/GR ratio, categorized as high (≥3, blue), moderate (2-<3, green), or low (<2, red), indicating the relative balance between the glutathione pool size and recycling capacity. Quadrants delineate distinct metabolic strategies, ranging from low-TKT reserve, non-oxidative bias (lower left), to high-TKT reserve, oxidative PPP bias (upper right). Axes represent relative prioritization among pathways rather than absolute enzymatic rates and are intended to summarize integrated metabolic allocation strategies across strains.

TEM exhibited markedly elevated G6PDH/basal TKT ratios, coupled with low (GSH+GSSG)/GR values, indicating a strong oxidative PPP bias with limited glutathione buffering capacity. In contrast, the LT strain had a much lower oxidative PPP bias and a comparatively large glutathione pool relative to recycling capacity. The ASN strain exhibited lower G6PDH/TKT ratios and moderately to highly elevated total glutathione/GR values, consistent with a greater reliance on non-oxidative PPP activity and a buffered redox capacity. TXD occupied an intermediate position, indicating a more balanced metabolic strategy along with the greatest consistency among individuals.

Separation of strains under identical ambient conditions within this three-variable metabolic space reflects distinct baseline organizational solutions to balancing carbon conservation, redox supply, and antioxidant resilience. This suggests that distinct brook trout strains exhibit contrasting biochemical allocation strategies that may contribute to their performance under differing environmental conditions, highlighting metabolic organization itself as an axis of physiological diversity.

### Metabolic organization in the context of strain diversity

The study fish were reared under standardized common hatchery conditions for eight months, minimizing environmental variation and allowing persistent strain-level differences in hepatic metabolism to be evaluated. Differences in liver protein allocation, redox pools, and PPP organization are consistent with intrinsic metabolic phenotypes reflecting their distinct strain-specific evolutionary and broodstock histories. Previous studies have documented persistent strain-specific differences in growth, thermal responses, survival and habitat use, with hatchery-introgressed populations frequently retaining watershed-specific genetic structure.[6, 25–30, 53, 63, 64] The coordinated differences observed here therefore extend these established phenotypic distinctions to the level of hepatic metabolic organization.

Variation in oxidative versus non-oxidative PPP capacity, together with differences in glutathione pool size relative to recycling capacity, indicates that strains employ alternative patterns of carbon allocation and redox metabolism. These constitutive metabolic phenotypes provide a physiological framework for understanding how genetic background may shape responses to environmental challenges. Given the escalating warming and low oxygen conditions in temperate lakes,[65] establishing these baseline metabolic organizations is an important foundation for interpreting strain-specific physiological responses under environmental stress.

## Conclusions

Brook trout strains reared under common ambient conditions differed fundamentally in hepatic metabolic organization, as reflected by hepatic soluble protein density, PPP enzyme capacity, glutathione metabolism, and ratiometric indices, exhibiting coordinated metabolic phenotypes that were not apparent from individual biochemical measurements. In particular, the marked divergence between the TEM strain and its domestic hybrid (TXD strain) illustrates how closely related strains can differ substantially in the organization of carbon metabolism and redox investment. These findings suggest that metabolic allocation represents a stable component of strain identity rather than a single optimal biochemical configuration. The framework presented here provides a broadly applicable approach for characterizing metabolic organization and establishes a foundation for understanding how constitutive metabolic phenotypes shape physiological responses to environmental stress. Future integration with environmental and physiological challenge studies will clarify how baseline metabolic organization across strains shapes adaptive performance under warming and hypoxic stress.

## Acknowledgements

We acknowledge the Adirondack Fishery Research Program (Cornell University) for donating the brook trout samples and the associated AFRP team (Pete McIntyre, Connor Reeve, and Evan Dlugos) for discussions, logistical assistance, and maintaining the fish. We thank Esther Angert for laboratory use at Cornell University.

## Contributions

KE supervised the laboratory work, analyzed the data, provided resources, and drafted the manuscript. ER designed the ambient temperature brook trout strain comparisons and collected and processed all liver samples. CK provided input to the fisheries application, access to the AFRP, and reviewed the manuscript. KE, ER, BM, and DK carried out tissue extractions and biochemical analyses.

## Data availability statement

The data supporting the findings of this study are available from the corresponding author upon reasonable request.

